# Dawn and photoperiod sensing by phytochrome A

**DOI:** 10.1101/253989

**Authors:** Daniel D Seaton, Gabriela Toledo-Ortiz, Akane Kubota, Ashwin Ganpudi, Takato Imaizumi, Karen J Halliday

## Abstract

In plants, light receptors play a pivotal role in photoperiod sensing, enabling them to track seasonal progression. Photoperiod sensing arises from an interaction between the plant’s endogenous circadian oscillator and external light cues. Here, we characterise the role of phytochrome A (phyA) in photoperiod sensing. Our meta-analysis of functional genomic datasets identified phyA as a principal transcriptional regulator of morning-activated genes, specifically in short photoperiods. We demonstrate that *PHYA* expression is under the direct control of the PHYTOCHROME INTERACTING FACTOR transcription factors, PIF4 and PIF5. As a result, phyA protein accumulates during the night, especially in short photoperiods. At dawn phyA activation by light results in a burst of gene expression, with consequences for anthocyanin accumulation. The combination of complex regulation of *PHYA* transcript and the unique molecular properties of phyA protein make this pathway a sensitive detector of both dawn and photoperiod.

**Significance statement:** The changing seasons subject plants to a variety of challenging environments. In order to deal with this, many plants have mechanisms for inferring the season by measuring the duration of daylight in a day. A number of well-known seasonal responses such as flowering are responsive to daylength or photoperiod. Here, we describe how the photoreceptor protein phytochrome A senses short photoperiods. This arises from its accumulation during long nights, as happens during winter, and subsequent activation by light at dawn. As a result of this response, the abundance of red anthocyanin pigments is increased in short photoperiods. Thus, we describe a mechanism underlying a novel seasonal phenotype in an important model plant species.

## Introduction

As photosynthetic organisms, plants are highly tuned to the external light environment. This exogenous control is exerted by photoreceptors, such as five member phytochrome family (phyA-E), that, in turn, regulate the activity of key transcription factors. An important feature of phytochrome signalling is that it can be strongly influenced by the plants internal circadian clock, which operates as a master regulator of rhythmic gene expression. The interplay between phytochrome signalling and the clock aligns daily gene expression profiles to shifts in day-length. These adjustments and associated post-transcriptional events form the basis of photoperiodic sensing, coordinating molecular, metabolic and developmental responses to the changing seasons.

Earlier work has shown that light and the clock interact through so called “external coincidence” mechanisms to deliver photoperiodic control of responses such as flowering time and seedling hypocotyl growth (1,2). Previously we used a modelling approach to assess the functional characteristics of these two external coincidence mechanisms (3). An important component of our study was the analysis of published genomics data that allowed us to identify new network properties and to test the applicability of our model to the broader transcriptome. This work highlighted the huge potential of data mining approaches to uncover new molecular mechanisms of external coincidence signalling.

A well characterised external coincidence mechanism involves the PHYTOCHROME INTERACTING FACTOR transcription factors PIF4 and PIF5, that regulate rhythmic seedling hypocotyl growth in response to short photoperiods. In this instance, sequential action of the clock Evening Complex (EC) and phyB defines the photoperiodic window during which PIF4/5 can accumulate. Light activated phyB is known to negatively regulate PIF4/5 by triggering their proteolysis and/or by sequestering PIFs from their target promoters (4,5). The EC, comprising EARLY FLOWERING 3 (ELF3), EARLY FLOWERING 4 (ELF4), and LUX ARRHYTHMO (LUX), is a transcriptional repressor that has a post-dusk peak of activity. In daily cycles that have a short night the EC completely blocks *PIF4/5* expression. In contrast, nights longer than 10-12h exceed the period of EC action, allowing *PIF4/5* to accumulate and regulate gene expression. The period of PIF activity is abruptly terminated at dawn, following activation of phyB by light. This external coincidence module therefore delivers a diurnal control of growth that is only active in short-day photocycles and becomes more robust as the night lengthens.

The diurnal PIF growth module provides a clear example of how phyB contributes to photoperiod sensing. The phytochrome family share a set core characteristics that enable tracking of light quality and quantity changes, such as those that occur at dawn. The phytochrome chromoproteins exist in two isomeric forms, inactive Pr and active Pfr, that absorb in the red (peak 666nm) and far-red light (peak 730nm), respectively. Red light drives photoconversion from Pr to Pfr, while far-red light reverses this process. This so called R/FR reversibility allows phytochromes to operate as biological light switches that respond to light availability spectral and quality. Once formed, the active Pfr translocates from the cytosol to the nucleus to perform its signalling functions.

The basic photochemistry of phytochrome signalling is conserved across the phytochrome family. However, phyA exhibits unique signalling features, including nuclear translocation kinetics and protein stability. As a result, the responses of phyA to light are distinctive. For example, phyB-E responses are classically R/FR reversible, while phyA responses are not. Instead, phyA is tuned to detect continuous FR-rich light, indicative of close vegetation, in the so-called far-red high irradiance responses (FR-HIR) (6). phyA also initiates very low fluence responses that are particularly important for activating germination and de-etiolation in low light scenarios (e.g. when shielded by soil, debris, or vegetation). Another distinguishing feature is that unlike phyB-E, that are light stable, the phyA holoprotein is unstable in the presence of light. These characteristics mean that in photoperiodic conditions phyA protein levels are robustly diurnal (7), though it is not clear what drives phyA re-accumulation during the night.

Considerable progress has been made in understanding the molecular mechanisms of phyA signalling (6). Upon exposure to R or FR light, phyA is activated and moves from the cytosol to the nucleus. Nuclear import requires the NLS-containing helper proteins FAR-RED ELONGATED HYPOCOTYL 1 (FHY1) and FHY1-like (FHL) (8). FHY1 and FHL shuttle back and forth between the nucleus and the cytosol, which is an important component of the FR-HIR (9). In the nucleus, phyA Pfr negatively regulates several proteins through direct interaction, including the PHYTOCHROME INTERACTING FACTOR (PIF) transcription factors, the E3 ligase component CONSTITUTIVE PHOTOMORPHOGENIC1 (COP1), and SUPPRESSOR OF PHYA-105 1-4 (SPA1-4) (10,11). The COP1/SPA complex targets several upstream transcription regulators, including LONG HYPOCOTYL 5 (HY5), LONG HYPOCOTYL IN FAR-RED 1 (HFR1), and LONG AFTER FAR-RED LIGHT 1 (LAF1), for degradation (12). Through the regulation of this suite of key transcription factors, phyA can modulate the expression of thousands of genes (13–15).

The activity of the phyA signalling pathway is regulated at multiple levels. The timing of *PHYA* expression is controlled by the circadian clock (16,17), and by light, though the underlying molecular mechanisms are currently unknown. phyA protein is both activated and destabilised by light (18). Thus, a full appreciation of phyA signalling can only be gained by studying the interplay between these layers of regulation. This can be achieved by analysing dynamics of phyA regulation and action through different photoperiods where the competing regulatory signals converge at different times. Previously we have constructed mathematical models to hone our understanding of photoperiodic control of flowering and PIF-mediated growth (3). The approach has been particularly useful for identifying non-intuitive pathway behaviours that arise from complex regulatory dynamics.

In this paper, we combine analysis of genome-scale datasets, mathematical modelling, and experimentation to unravel the molecular mechanisms of phyA regulation in light/dark cycles. We show that *PHYA* is directly targeted by the transcription factors PIF4 and PIF5. These transcription factors are under the dual control of light (via phytochromes (4)) and the circadian clock (via the evening complex (19)). This regulation results in dynamic regulation of *PHYA* transcript abundance, leading to high accumulation at night in short photoperiods. At dawn, phyA then induces the expression of hundreds of genes, including genes involved in anthocyanin biosynthesis. This firmly establishes a role for phyA as a sensor of dawn and short photoperiods.

## Results

### Data mining identifies phyA as a potential short-photoperiod sensor

Our previous work applied data mining methods to derive new molecular understanding of light signalling (3). In this study we used data mining to identify gene regulatory mechanisms that respond to changing photoperiod. This approach was made possible by the high quality transcriptomic and ChIP data for diurnal and light-controlled gene expression (Table S1; Table S2). To do this we developed a computational workflow combining co-expression clustering and gene set enrichment (Fig 1A). First, genes were clustered on the basis of expression in a variety of conditions, focussing on different light conditions, and mutants of circadian and light signalling pathways (see Table S1 for a complete description of datasets). Importantly, this included gene expression in long photoperiods (16h light: 8h dark (8L:16D) and short photoperiods (16L:8D). This procedure identified 101 co-expression clusters (Datafile 1).

**Figure 1.**
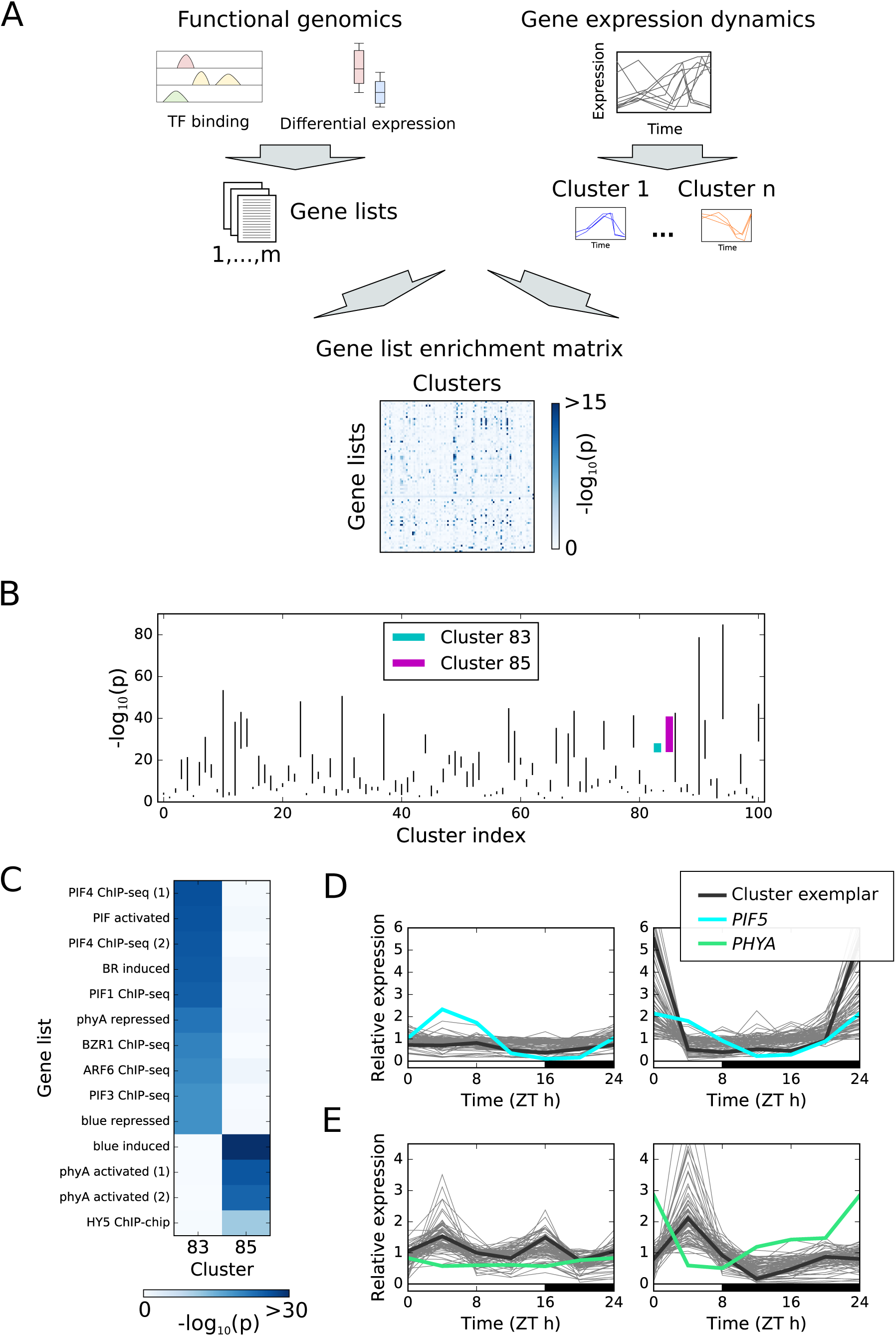
Mining functional genomic data for active gene regulatory networks. (A) Flowchart of data integration. Genes were clustered together according to their dynamics in a range of conditions. Functional genomic datasets (e.g. ChIP-seq, RNA-seq) were curated from literature in the form of gene lists. Each cluster was then tested for over-enrichment of each gene list (hypergeometric test). (B) Top gene list enrichment scores across all clusters. Vertical lines indicate the range spanned by the three top-scoring enrichments. (C) Highlighted enrichment tests for clusters 83 and 85, which are enriched for distinct subsets of phytochrome-related gene lists. (D) Short day, night-specific expression of cluster 83, and its relationship with *PIF5* expression. (E) Short day, morning-specific expression of cluster 85, and its relationship with *PHYA* expression.

To increase the likelihood of identifying regulatory mechanisms, we assessed a broad range of potential regulatory pathways. To do this, we consolidated 527 gene lists from available datasets. This consisted of 140 gene lists from 47 papers, covering a broad range of regulatory pathways (e.g. responses to hormones, response to stimuli, ChIP-seq of transcription factors; see Table S2 for descriptions), combined with a further 387 transcription factor binding datasets generated in high throughput by DNA affinity purification sequencing (DAP-seq) (20). For each cluster of co-expressed genes, if there is a significant overlap between a particular gene set (e.g. differential expression or transcription factor binding) and the genes in a particular cluster, it can suggest potential regulatory mechanisms. Here, enrichment is quantified by the p-value of overlap between gene sets and clusters (hypergeometric test; see Table S3 for all calculated values). Similar approaches have previously been used to identify gene regulatory networks in a wide variety of contexts (e.g. (21–23)). Analogous approaches include the identification of promoter motifs by enrichment in give gene sets (e.g. (24)). We have developed a simple software tool, AtEnrich (“Arabidopsis thaliana gene list Enrichment analysis”), for performing combined clustering and enrichment analysis of these gene lists (Datafile 2).

Enrichment analysis identified many high-scoring associations, with 37 of 101 clusters enriched with at least one gene set at p < 10^−20^ (Fig 1B). As expected, this approach highlighted roles for circadian and light signalling factors in controlling the diurnal dynamics of gene expression. In particular, phytochrome signalling is prominently implicated in the regulation of several clusters. One example of this is Cluster 83, which is regulated by the *PIF4/PIF5* pathway, that controls changes in hypocotyl elongation with photoperiod (3,25) (Fig1C,D). Targets of the PIF family of transcription factors (also including PIF1 and PIF3) have been identified by ChIP-seq (26–28), as have targets of PIF-interacting proteins including AUXIN RESPONSE FACTOR 6 (ARF6) and BRASSINAZOLE-RESISTANT 1 (BZR1) (29). Cluster 83 is strongly enriched for all of these gene lists (p<10^18^ in all cases; hypergeometric test; Fig 1C). PIF4 and PIF5 are known to be abundant and active in the dark. Due to transcriptional repression by the clock repressor EC, they accumulate during the night specifically in short photoperiods, and in the *lux* mutant (which lacks the EC) and *LHY* overexpressor (which has reduced EC activity) (3,25,30). The expression profile of cluster 83 genes in long days (16L:8D) and short days (8L:16D) is consistent with this understanding of the pathway. This is illustrated in Fig 1D, with higher night-time levels of *PIF5* transcript in short photoperiods, and higher night-time expression of genes in this cluster. As expected, this cluster includes well-known markers of PIF activity including *ATHB2*, *IAA29*, *HFR1*, and *CKX5* (30).

Phytochrome signalling, and in particular phytochrome A, is also implicated in the regulation of cluster 85. Analysis revealed that this cluster is enriched for genes responding rapidly to red light in a phyA-dependent manner (14), and genes responding to far red light in a phyA-dependent manner (13) (Fig 1C). Furthermore, it is enriched for genes bound by the transcription factor HY5 (31), which is stabilised by phyA via its interaction with COP1 (32). This cluster of genes also displays a pattern of gene expression consistent with sensitivity to light, with a peak in expression following dawn (Fig 1E). The size of this peak changes with photoperiod, and is especially pronounced in short photoperiods (Fig 1E). Interestingly, the expression of the genes in the morning is correlated with expression of *PHYA* during the preceding night, which is higher during the night in short photoperiods (Fig 1E). Therefore, we proceeded to investigate the photoperiodic regulation of *PHYA* expression, and the implications of this for the seasonal control of gene expression of this set of genes.

### A model of PIF activity predicts *PHYA* expression dynamics

Previous reports have indicated that phyA protein accumulates in etiolated seedlings and during the night in a diurnal cycle through an unknown process (7,33). As highlighted by earlier studies and our clustering analysis, the PIF family of transcription factors display a similar pattern of activity (1,3,25). Furthermore, our previous analysis of gene expression dynamics identified *PHYA* as a putative target of PIF4 and PIF5 (3).

In order to assess the plausibility of the hypothesised regulation of *PHYA* expression by PIF4/5, we tested whether our model of PIF4/5 activity was able to explain *PHYA* dynamics in different photoperiods and circadian clock mutants. This model is presented schematically in Fig 2A. In short days (8L:16D), both model and data exhibit rhythmic *PHYA* expression with an end of night peak (Fig 2B). In long days (16L:8D), however, expression is low throughout the day and night (Fig 2B). The model also matches the measured response of *PHYA* expression at end of night and end of day across multiple photoperiods (Fig S1). Finally, the model matches the exaggerated nocturnal rise in *PHYA* observed in two circadian clock mutants - the *lux* mutant and *LHY* overexpressor (Fig 2C,D). These mutants are notable for exhibiting weak evening complex activity, with a resultant increase in *PIF4* and *PIF5* expression during the night. In summary, a model of PIF4/5 regulation of *PHYA* is able to explain differences in *PHYA* expression across a range of environmental conditions and genotypes.

**Figure 2.**
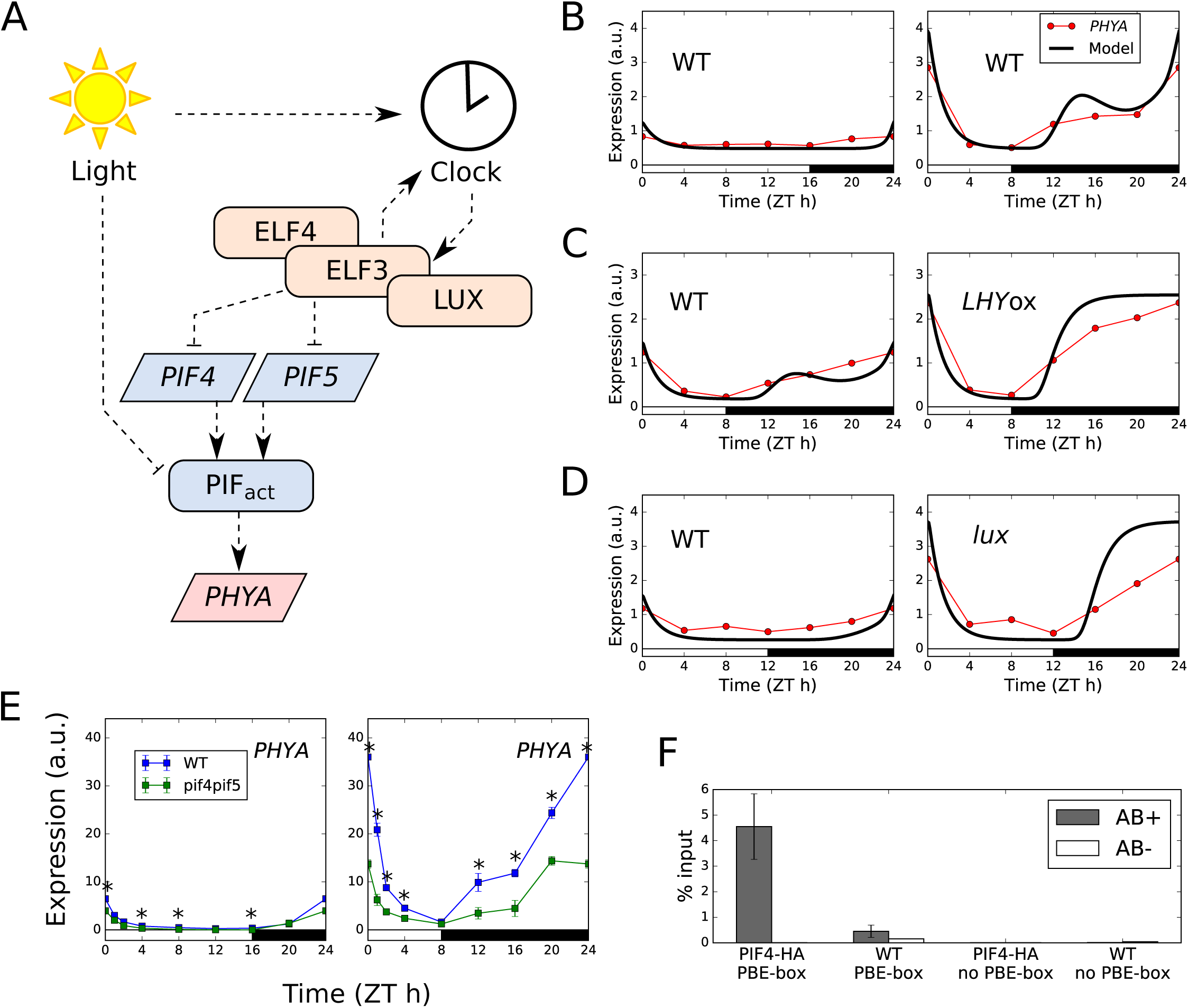
*PHYA* expression is directly regulated by PIF4 and PIF5. (A) Schematic of a model of PIF signalling, extended to include regulation of PHYA (Seaton et al, 2015). (B-D) Comparison of model simulations and microarray data for *PHYA* in short compared to long photoperiods (B), WT (Ler) compared to *LHY*ox in 8L:16D light/dark cycles (C), and WT (Col) compared to the *lux* mutant in 12L:12D light/dark cycles (D). (E) *PHYA* expression in short and long photoperiods, in the WT (Col) and the *pif4;pif5* mutant. Plants were grown for 2 weeks in the given photoperiod. Expression was measured relative to *ACT7*. (n=3, error bars represent SEM, ZT0 timepoint re-plotted at ZT24). (F) ChIP-qPCR of PIF4 binding to the *PHYA* promoter. Plants were grown for two weeks in short days (8L: 16D white light, 100 µmol/m^2^/s) at 22°C, and samples were collected at the end of the two weeks at ZT0 (n=3, error bars represent SEM).

Further support for the regulation of *PHYA* by PIF4/5 comes from existing microarray and RNA-seq datasets. These show that *PHYA* levels are reduced in etiolated and shade-grown seedlings lacking PIF1, PIF3, PIF4, and PIF5 (i.e. the *pifQ* mutant (*pif1*;*pif3*;*pif4*;*pif5*)) (34,35) (Fig S2). Interestingly, the *PHYA* cofactor *FHL* (also identified as a possible PIF4/5 target in (3)) shows similar patterns of expression across the range of microarray datasets inspected here, and its expression can also be explained by the model of PIF4/5 activity (Figs S2, S3). This suggests that PIF4/5 regulate both *PHYA* and *FHL*, and therefore may exert significant influence on the activity of the phyA signalling pathway.

### PIF4 and PIF5 directly regulate *PHYA* expression

To further establish a role for PIF4 and PIF5 in regulating *PHYA* and *FHL* expression, we measured mRNA levels by qPCR in Col0 (wild type) and *pif4;pif5* plants, in short (8L:16D) and long (16L:8D) photoperiods. This revealed the expected *PHYA* expression profile, with transcript levels rising to much higher levels during the night in a short day compared to a in a long day. *PHYA* expression was markedly reduced in the *pif4;pif5* mutant specifically in short photoperiods (Fig 2E) and was reduced further in the *pifQ* mutant, that lacks PIF1 and PIF3 in addition to PIF4 and PIF5 (Fig S4). Furthermore, a similar pattern was observed for *FHL*, as expected (Fig S4). These data further implicate PIFs in regulation of *PHYA* and *FHL*. As for transcript, phyA protein accumulates to higher levels in short days compared to long days (Fig S5A), and its levels at ZT0 in short days are reduced in the *pif4;pif5* and *pifQ* mutants (Fig S5B). These data suggest that PIFs may act collectively to regulate phyA abundance.

The strong coordination between *PHYA* expression and PIF activity across many conditions (i.e. different photoperiods, *pif* mutants, and clock mutants) suggested that this regulation might be direct. Numerous ChIP-seq analyses of the PIF family have been performed across a range of conditions (e.g. in deetiolated seedlings (27,28,36) and in low R:FR ratio conditions (26)). Among these, only (36) has found direct binding of a PIF (PIF4) to the *PHYA* promoter, in deetiolated seedlings. In order to test direct regulation of *PHYA* by PIFs in our conditions, we performed ChIP for PIF4-HA and PIF5-HA on the *PHYA* promoter in plants grown in short days, focussing on a region with a PIF-binding E-box (PBE) element (CACATG; (28)). The results of this are shown in Fig 2F (PIF4) and Fig S6 (PIF5), with enrichment of PIF4-HA and PIF5-HA at the *PHYA* promoter. Thus, PIF4 and PIF5 appear to regulate *PHYA* expression by direct binding to its promoter in short days.

### PIFs regulate phyA action specifically in SDs

Additional support for PIF4 and PIF5 as SD regulators of *PHYA* comes from our hypocotyl data. When supplied continuously, far-red light activates phyA in an HIR mode (18). We used this unique photochemical property to provide a readout for phyA presence through the night of SD- and LD-grown seedlings. Our data show that 4h of FR light (delivered at the end of the night) suppresses hypocotyl elongation in a phyA and PIF-dependent manner in SDs but not LDs (Fig S7). To rule out any potential influence of phyB and other light stable phytochromes on phyA action we also provided brief end-of-day (EOD) far-red treatments that switch these phytochromes to their inactive Pr conformer. As expected, EOD deactivation of phyB enhanced hypocotyl elongation in WT and *phyA* seedlings, and this was more marked in SDs. Delivery of prolonged far-red to EOD-far-red treated seedlings, again led to phyA-suppression of hypocotyl elongation, a response that was markedly reduced in *pif4:pif5* and *pifQ* mutants. These photo-physiological experiments provide robust support for our central hypothesis that the photoperiodic phyA regulation is largely conferred by SD PIF action.

### phyA mediates a photoperiod-dependent acute light response

Differences in phyA accumulation during the night are expected to result in differences in phyA activity during the following day. In order to assess this, we developed a model of phyA signalling mechanisms, as a simplified version of the model of Rausenberger et al. (9) (Fig 3A). In this model, phyA signalling activity is high when light is present and phyA protein is abundant. The rapid decrease in the level of phyA protein after dawn means that phyA activity peaks in the early morning regardless of conditions. This pulse is termed an 'acute light response'. This is illustrated in Fig 3B, showing simulations of the combined clock-PIF-phyA model in short and long photoperiods.

**Figure 3.**
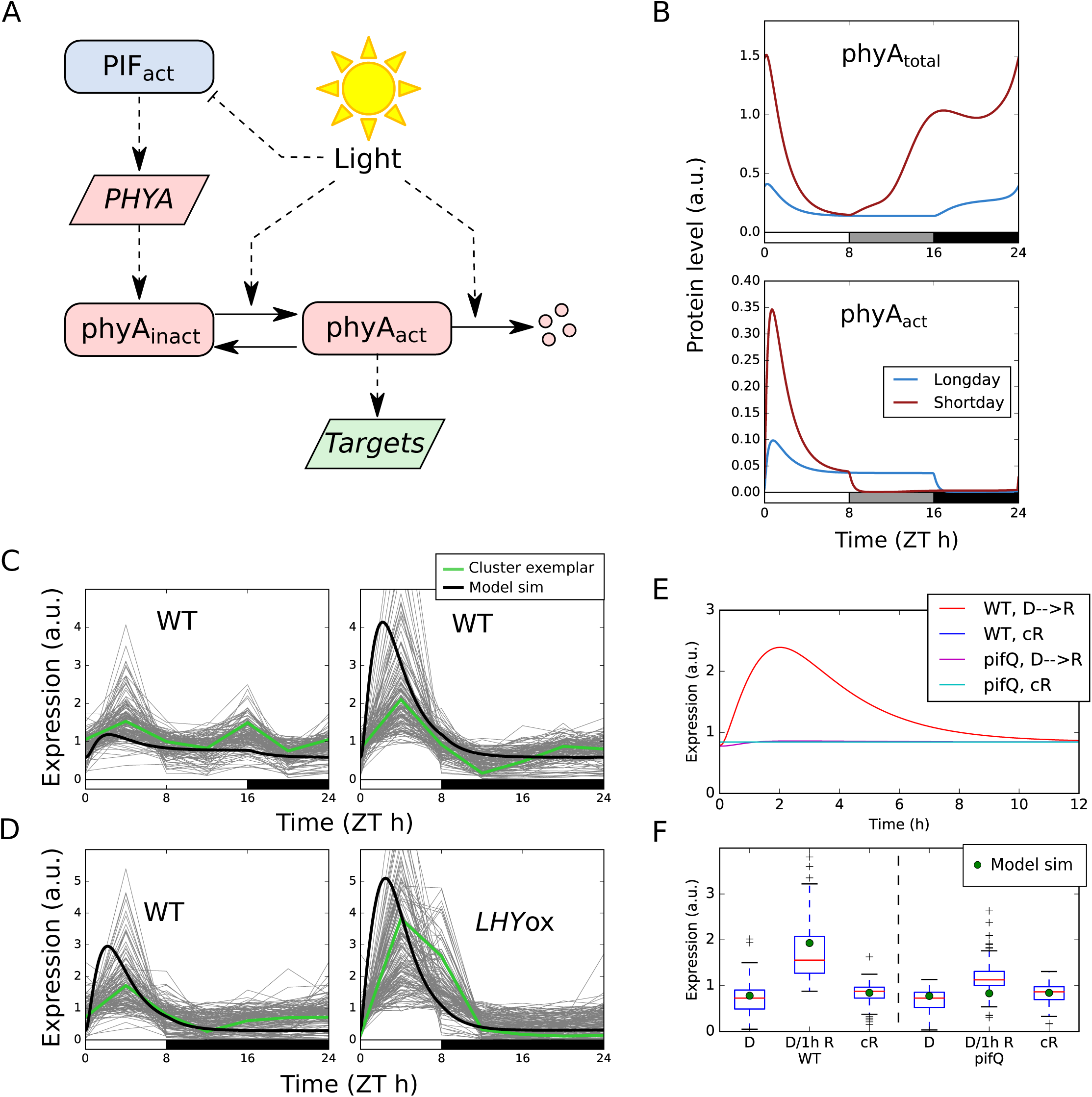
A model of phyA signalling predicts gene expression dynamics. (A) Model schematic. Solid lines represent mass transfer, dashed lines represent regulatory effects. Transcripts are represented by trapezoids, proteins by rectangles. (B) Simulation of a simple model of phyA signalling in short and long photoperiods. (C, D) Gene expression of the putative phyA-regulated cluster of co-expressed genes, compared to model simulations, in photoperiods (C), and LHYox (D) (data from Michael et al. 2008). (E) Simulated expression of a putative phyA target, following a transition from darkness to continuous red light. (F) Gene expression of the putative phyA-regulated cluster of co-expressed genes following a transition from darkness to continuous red light (data from Leivar et al, 2009).

The model predicts that the changing activity of PIFs across different photoperiods and genotypes changes the amplitude of the acute light response (Fig 3B). In particular, it predicts that the amplitude of the acute light response at dawn is increased in short photoperiods, as well as in the *LHYox* line and the *lux* mutant (i.e. conditions with high *PHYA* expression during the night). The genes in the putative phyA-regulated cluster (cluster 85) display these dynamics (Fig 3 C,D). The model is also able to make predictions for gene expression dynamics during seedling deetiolation, in which dark-grown seedlings are exposed to red light. Here, the model predicts a diminished amplitude of response in the *pifQ* mutant during deetiolation in red light (Fig 3E). Again, genes in cluster 85 display dynamics consistent with the model across these conditions when compared to a microarray dataset in which plants were grown in darkness and treated with red light for 1h, or grown in continuous red light (35) (Fig 3F). Together, these results demonstrate that our molecular understanding of this pathway is consistent with phyA regulation of cluster 85, as expected based on its enrichment for phyA-associated terms in our meta-analysis of functional genomic datasets (Fig 1C).

In summary, this cluster of putative phyA targets displays expression dynamics consistent with our mechanistic understanding of phyA signalling, as captured by our mathematical model. This further implicates phyA as a key regulator of these genes.

### phyA is a clock input specifically in short photoperiods

PhyA is known to mediate light input to the circadian clock (2,37,38). This suggests that PIF4 and PIF5 could play a role in the clock, through their regulation of phyA. One candidate target for phyA signalling in the circadian clock is the dawn-induced *PSEUDO RESPONSE REGULATOR 9* (*PRR9*). The induction of *PRR9* after dawn is very sensitive to photoperiod, with a strong induction in short photoperiods (39). This pattern is not explained by current models of the circadian clock, suggesting the existence of an unknown regulatory mechanism (39). Furthermore, the *PRR9* promoter has been identified as bound by phyA, HY5, and FHY1 (13,31,40). Measurement of *PRR9* expression in *pif4;pif5* and *phyA* demonstrates that *PRR9* is indeed regulated by phyA, with reduced expression in both mutants (Fig S8A). This is consistent with molecular data from ChIP-seq experiments showing that *PRR9* is bound by HY5, FHY1, and phyA (13,31,40). As expected, this difference is specific to short photoperiods.

Given the significant effect of phyA on *PRR9* expression, we hypothesised that this regulation could act as a rephasing mechanism for the clock. To test this hypothesis we assayed the expression of the core clock genes *PRR7*, *TOC1*, *GI*, *LUX*, and *ELF4* in *phyA* and *pif4;pif5* mutants in short and long days (Fig S8B). While statistically significant differences between WT and both *phyA* and *pif4;pif5* are observed for most transcripts at several timepoints, the fold-changes in gene expression were generally modest (Fig S8B). This suggests that the short day component of phyA action does not have a strong effect on circadian clock dynamics in our conditions.

These results are consistent with a previous report that loss and overexpression of PIFs has little effect on clock gene expression in general (1,41). In summary, our results demonstrate that phyA regulates *PRR9* specifically in short photoperiods.

### phyA confers photoperiodic control of anthocyanin accumulation

Our results demonstrate that phyA-mediated acute light responses are amplified in short photoperiods. Therefore, we expect short photoperiods to exaggerate *phyA* mutant phenotypes. In order to identify potential phenotypes of interest, we assessed enrichment of gene ontology (GO) terms within the cluster of putative phyA targets. This identified highly significant enrichment for antocynanin and flavonoid biosynthesis (GO:0046283, GO:0009812; Table S4). This is consistent with the observation that phyA is involved in anthocyanin accumulation in far-red light (42,43), and regulates expression of *CHALCONE SYNTHASE* (*CHS*), an enzyme involved in the synthesis of flavonoid and anthocyanin precursors.

To test the phyA photoperiodic link, we measured expression of *FLAVANONE 3-HYDROXYLASE* (*F3H*) and *CHS* in short and long days, in WT (Col0), *pif4;pif5*, and *phyA*. Although *CHS* was not identified in the phyA-regulated cluster (cluster 85), it is a well-known target of phyA signalling, and displays several of the expected features of induction by phyA in available microarray data, including a photoperiod-modulated dawn peak. Our timeseries qPCR data show that in short days *CHS* and *F3H* transcript levels rise rapidly post-dawn in WT, but this response is markedly reduced in *phyA* and *pif4;pif5* (Fig 4A). Contrasting with this, expression of *CHS* and *F3H* is similar in *phyA* and *pif4;pif5* through a long day (Fig 4A). These data are consistent with phyA being most active during the day in short photoperiods.

**Figure 4.**
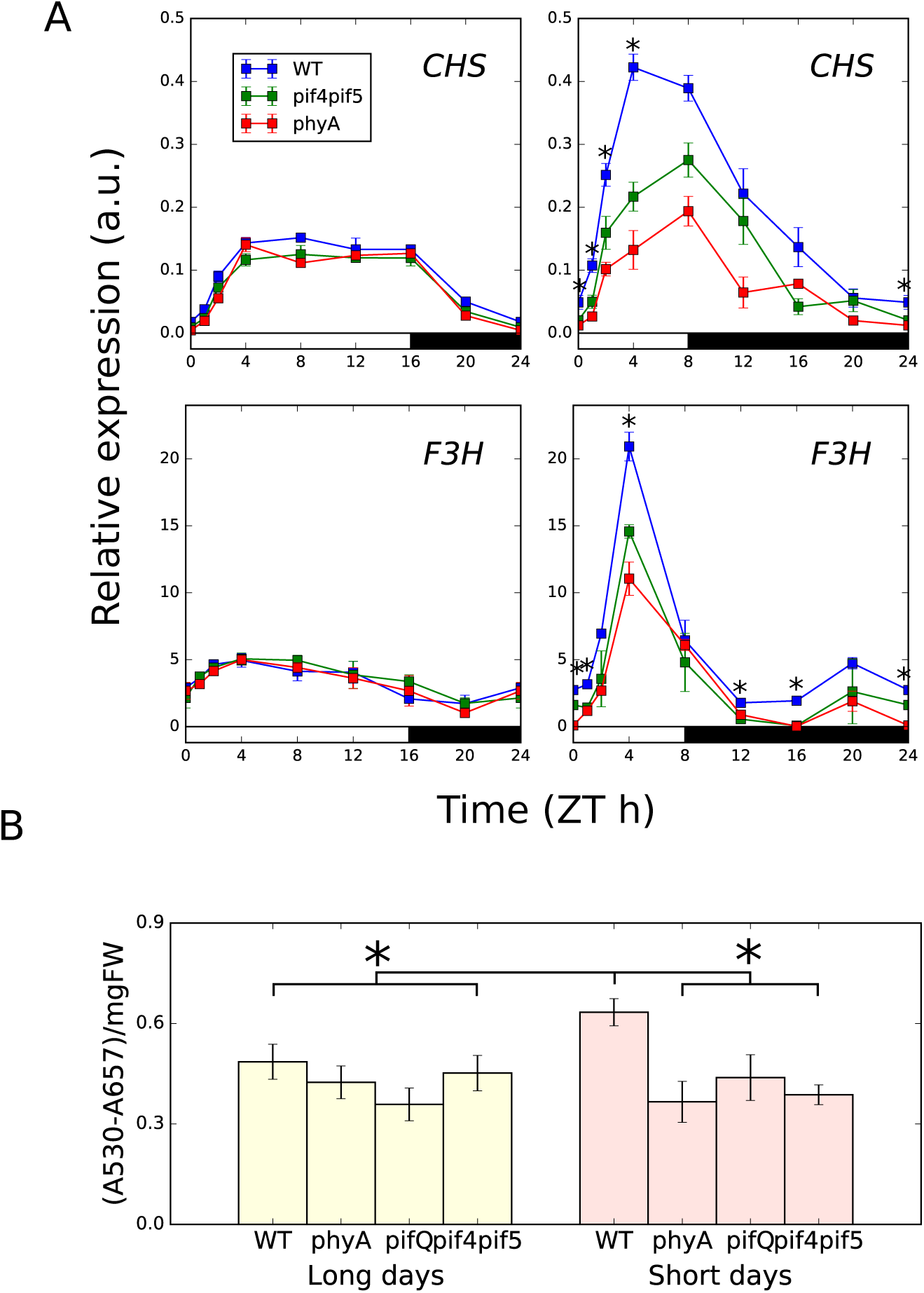
Anthocyanin accumulation is regulated by phyA in a photoperiod-specific manner. (A) qPCR timecourse data for F3H and CHS in short and long photoperiods, in WT (Col0), pif4;pif5, and phyA. Expression is relative to ACT7. Plants were grown for 2 weeks at 22°C under 100 µmol/m2/s white light in the specified photoperiod (* indicates significant difference at p<0.05 between WT and both pif4;pif5 and phyA, two-tailed t-test, n = 3, error bars represent SEM) (B) Anthocyanin accumulation in the same conditions as (A), also including the pifQ mutant. (* indicates difference from WT in short days at p < 0.01, one-tailed t-test, n = 3, error bars represent SD).

In order to test whether these differences in gene expression result in differences in metabolic phenotype, we measured anthocyanin accumulation in plants grown in short and long days. As expected, anthocyanin levels are highest in the WT in short days, and are reduced in the *phyA*, *pif4;pif5* and *pifQ* mutants, specifically in short days (Fig 4B). These results highlight a role for the PIF-phyA module in mediating seasonal changes in anthocyanin levels.

### phyA is a robust sensor of natural dawns

In the preceding experiments, the photoperiods applied included only two light levels: on and off. However, in the natural environment fluence rate increases gradually following dawn. To test whether the acute light responses we observed were the result of the binary on/off photoperiods applied, we measured *PHYA, F3H* and *CHS* expression across dawn in three variations of the short photoperiod: instantaneous dawn (i.e. the on/off light condition applied previously), fast dawn (reaching 100 µmol m^−2^ s^−1^ after 50min), and slow dawn (reaching 100 µmol m^−2^ s^−1^ after 90min). The timing of fast dawn was based on weather data from Edinburgh, UK, in short photoperiods (Fig S9; Supporting Information). First, the post-dawn *PHYA* mRNA profile was very comparable in WT and *pif4:pif5*, indicating consistent PIF4/5 control of PHYA across the different dawns. While the amplitude varied slightly, the expression profiles of *F3H* and *CHS* in WT, *phyA*, *pif4;pif5* and *phyA;pif4;pif5* were qualitatively similar in abrupt, fast and slow dawns (Fig S9). For the phyA target genes *F3H* and *CHS*, the impact of the *phyA;pif4;pif5 triple* mutant was more marked than monogenic *phyA* allele at ZT4 indicating that PIF4/5 partly operate independently of phyA at this time point. Collectively, these data show that the phyA-mediated acute response is maintained in simulated natural dawns where light levels ramp-up slowly or rapidly. This response consistency most likely results from inherent photosensory properties that enable phyA to detect and react to very low fluence rate dawn light.

## Discussion

Perception of light allows plants to prepare for the predictable daily and seasonal rhythms of the natural environment. We have delineated a role for the light photoreceptor phyA in both daily and seasonal responses. On a daily timescale, phyA acts as a precise sensor of dawn, peaking in activity following first light. On a seasonal timescale, the amplitude of this dawn peak in activity changes, and is especially pronounced in short photoperiods.

The ability of phyA to respond sensitively to dawn relies on two key properties: its ability to sense very low levels of light (44), and its accumulation in darkness (7,33). It is well established that the active Pfr form of phyA is light labile, and degrades fairly rapidly following light exposure. However, inactive phyAPr accumulates in seedlings that are kept in prolonged periods of darkness (7). A night-time rise in phyA protein levels has also been reported for seedlings grown in short days (33). Here, we have identified the PIF transcription factors as regulators of this nocturnal elevation in phyA, and linked this accumulation to the induction of hundreds of transcripts at dawn.

This cycle of accumulation and repression of photosensitivity across a dark-to-light transition is reminiscent of responses in the mammalian eye. A combination of physiological and molecular mechanisms heighten photosensitivity during prolonged darkness, but this sensitivity gradually diminishes during prolonged exposure to light (45). Such systems have been shown to enable sensitive responses to fold-changes in stimuli (46). This may be especially important in the case of phyA, as it allows a high-amplitude response at dawn, when there is a transition from darkness to low-intensity light. Furthermore, phyA is not the only light-labile photoreceptor: Cryptochrome 2 shows similar patterns of accumulation in darkness (33,47). Thus, our analysis of phyA signalling may have implications for other light signalling pathways. In particular, it highlights the importance of studying such pathways in conditions that approximate the natural environment i.e. in photoperiods.

Our analysis suggests that nocturnal accumulation of phyA results in photoperiodic responses. In short photoperiods, higher levels of phyA are present during the night, leading to an enhanced sensitivity to light at dawn. Inspection of transcriptomic and functional genomic datasets revealed that this expectation is met in hundreds of phyA-induced genes. Furthermore, these changes in gene expression have consequences for plant metabolism and growth. For example, induction of genes involved in flavonoid and anthocyanin biosynthesis in short photoperiods is reflected in changes in anthocyanin accumulation in these conditions. A role for phyA in regulating anthocyanin metabolism has previously been demonstrated under far red light (43). Here, we extend this role to plants grown under white light in short photoperiods.

The potential relevance of increased anthocyanin accumulation to growth in short photoperiods remains to be understood, but may involve protection from abiotic stresses (48). This establishes a novel mechanism and role for phyA in photoperiod responses, in addition to its previously described role in photoperiodic flowering (49).

Another interesting observation is role of the PIF-phyA module in controlling the expression of the clock component *PRR9*. While a role for the PIF family of transcription factors in regulating the circadian clock has long been suspected (50), our results represent the first demonstration of a mechanism for this regulation, through the control of the clock transcription factor PRR9. This constitutes a potential feedback loop in the circadian clock, a possibility first highlighted by studies demonstrating circadian control of *PHYA* transcription (16). While a strong effect of this pathway on expression of clock genes besides *PRR9* was not observed in our conditions, other conditions may produce a stronger effect. In particular, phyA is expected to be especially active in FR-rich vegetation shade, as occurs commonly in nature. A role for the PIF-phyA module in regulation of the clock under shade conditions remains to be ascertained.

Previously, substantial focus has been placed on the role of phyA in seedling establishment (18). We recently demonstrated a role for phyA, alongside other phytochromes, in resource management and biomass production (51), while others have shown that phyA contributes to the photoperiodic flowering response (49). Our study firmly positions phyA as a photoperiodic dawn sensor that is tuned to detect the very low light levels that signify dawn onset in the natural environment. This property ensures that phyA is a very reliable sensor of dawn transition in nature, as the weather, local and seasonal change can profoundly affect the intensity of morning light.

## Materials and Methods

### Coexpression clustering

The gene expression datasets used for clustering were microarray timeseries from short days vs long days in WT (Ler) (24), WT (Col) vs lux (24), WT (Ler) vs LHYox (24), and WT (Col) vs *pifQ (35)*, and RNA-seq timeseries from WT (Col) vs lnk1;lnk2 (52). See Supporting Information for details of the clustering method and similarity metric.

### Plant material and growth conditions

Columbia-0 (Col-0) wild type and mutants were used for experiments. The mutant alleles corresponded to: *pif4pif5* (*pif4-2*, *pif5-2*), *pifQ* (*pif1-1*, *pif3-3*, *pif4-2*, *pif5-2*). Over expressing plants included 35S::PIF4-HA and 35S::PIF5-HA. All have been previously described (26,34). Seeds were surface sterilized, sown in GM-agar media and stratified in darkness for 3 days at 4°C before given a 3 h white light pulse to induce germination. Seedlings were kept in the dark for 2 days at 22°C and transferred to Short Days (8L:16D) or Long Days (16L:8D) (22°C, white light 100 µmol m^−2^ s^−1^) for two weeks before harvesting at the indicated time. All samples were processed in biological triplicates.

### RNA isolation and transcript levels analysis by qPCR

For quantitative PCR (qPCR) experiments seedlings were prepared and sown as previously described (plant material and growth conditions above; see Supporting Information for details).

### Chromatin Immunoprecipitation assays

ChIP assays were conducted according to (53) except that 2 week old plants were used for the assay. Plants were grown for two weeks in short days (8L:16D white light, 100 *µ*mol/m^2^/s) at 22°C, and samples were collected at the end of the two weeks at ZT0. The sequences of the primers used in these experiments to amplify PBE-box containing promoter region of phyA are shown in Table S5.

### phyA Immunoblots

Total proteins were extracted from 100 mg of tissue from plants grown under short or long days for two weeks (see plant material and growth conditions) and harvested at the indicated times on day 14. Two separate experiments were performed for Fig S5A and B (see Supporting Information for details).

### Mathematical model of phyA signalling

The model of phyA signalling is an ODE model based on a simplification of the model by Rausenberger et al (9), integrated with ODE models of the circadian clock and PIF signalling pathways (3,54). A schematic is shown in Fig S10. Here, the model equations are presented and justified in detail. Parameter values are provided in Table S6.

*PHYA* transcript is governed by the equation:

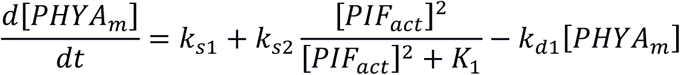

phyA protein in the inactive (R) form is then given by:

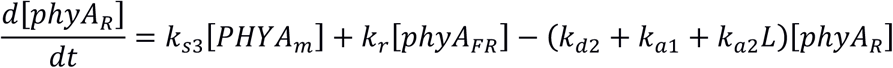

phyA protein in the active (FR) form is given by:

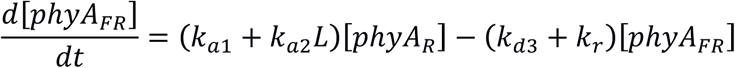

Levels of a downstream transcript are then given by:

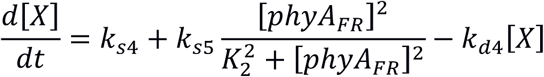

## Supplementary Information

**Table S1, gene expression dataset descriptions.** List of datasets used for clustering genes based on co-expression.

**Table S2, gene list descriptions.** Short descriptions of all curated gene lists taken from literature.

**Table S3, cluster enrichment scores.** -log10(pval) for the overenrichment of each gene list (rows) in each cluster (columns).

**Table S4, cluster 85 GO enrichment.** Top-scoring GO enriched terms of the genes in cluster 85.

**Table S5, primers.** PCR primer sequences for qPCR and ChIP-PCR analyses.

**Table S6, model parameters.** Parameters for the ODE model of phyA signalling.

**Datafile S1, gene clustering.** Tab-separated file listing gene IDs (left-hand column) and their corresponding cluster (right-hand column).

**Datafile S2, AtEnrich.** Tarzipped folder containing AtEnrich software for performing gene list and cluster enrichment analyses. Install from the command line with ‘python setup.py install’.

**Datafile S3, Gene list files.** A collection of gene lists collected from literature, that are analysed by AtEnrich.

## Acknowledgements

This work was partly supported by BBSRC grants (BB/M025551/1 and BB/N005147/1) to K.J.H., by NIH grants (GM079712) and Next-Generation BioGreen 21 Program (SSAC, PJ011175, Rural Development Administration, Republic of Korea) to T.I., and by Royal Society grant (RG2016R1) to G.T.. A.K. is supported by JSPS Postdoctoral Fellowships for Research Abroad. We thank James Furniss for assistance with plant growth, and members of the Halliday lab for comments on the manuscript.

---

**Supplementary Figure S1.**
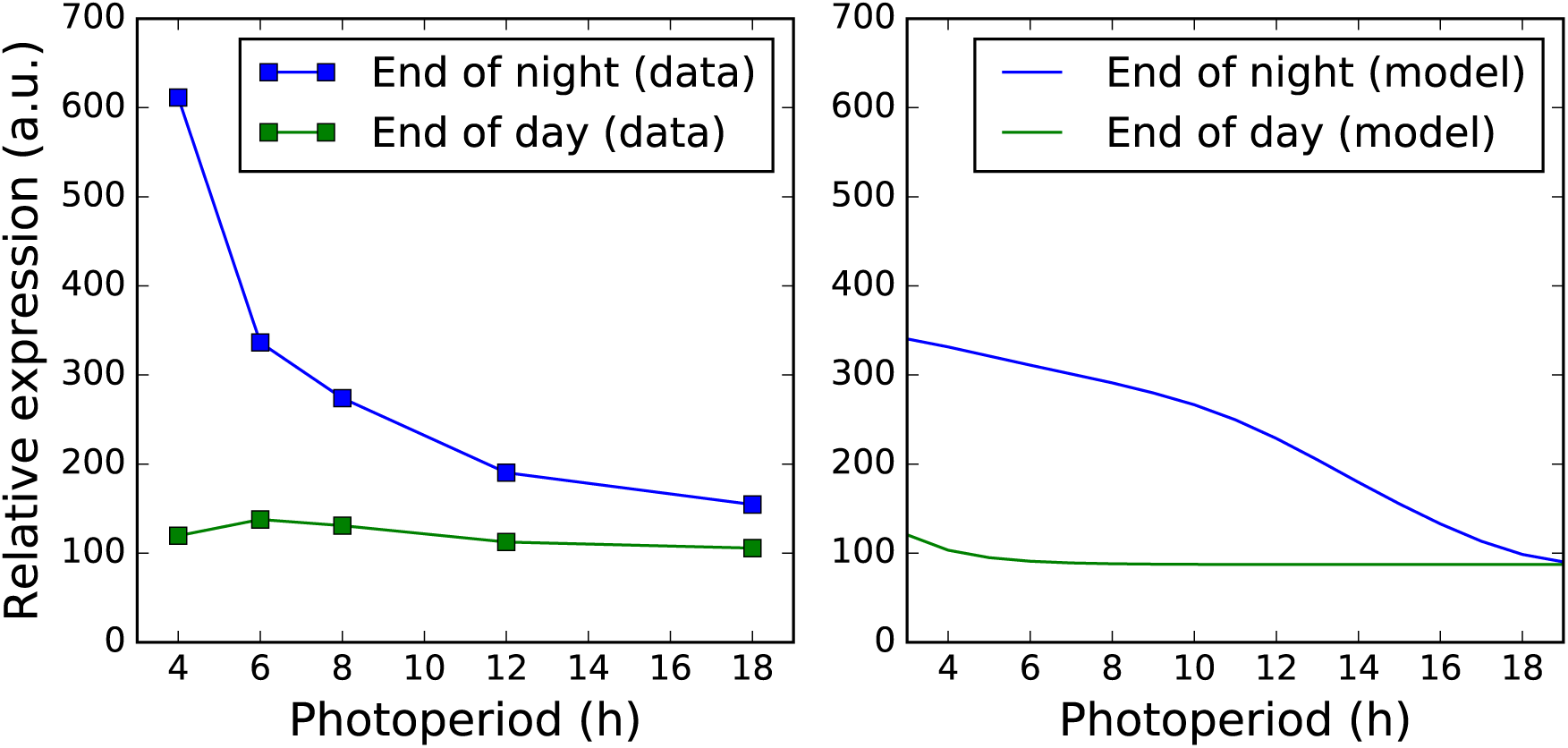
Comparison of model simulations and microarray data for *PHYA* expression at the end of night and end of day across 5 photoperiods (data from Flis et al, 2016).

**Supplementary Figure S2.**
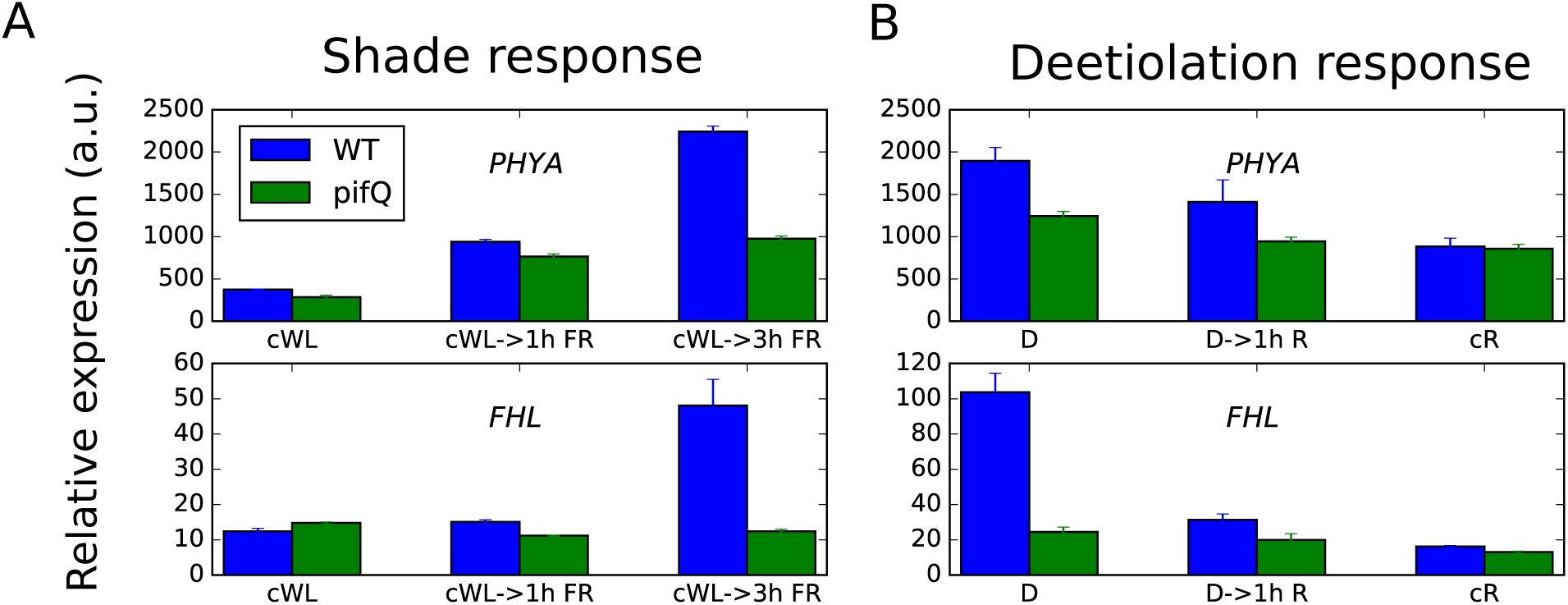
Expression of *PHYA* and *FHL* in response to shade and deetiolation. (A) Shade response microarray data are from Leivar et al, 2012. WT and pifQ seedlings were grown for 2 days in white light (cWL), supplemented by far red light for 1 h (cWL-> 1h R), or supplemented by far red light for 3h (cWL->3h FR). (B) Deetilation response microarray data are from Leivar et al, 2009. WT and *pifQ* seedlings were grown for 2 days in the dark (D), followed by 1 h in red light (D-> 1h R), or grown for 2 days in red light (cR).

**Supplementary Figure S3.**
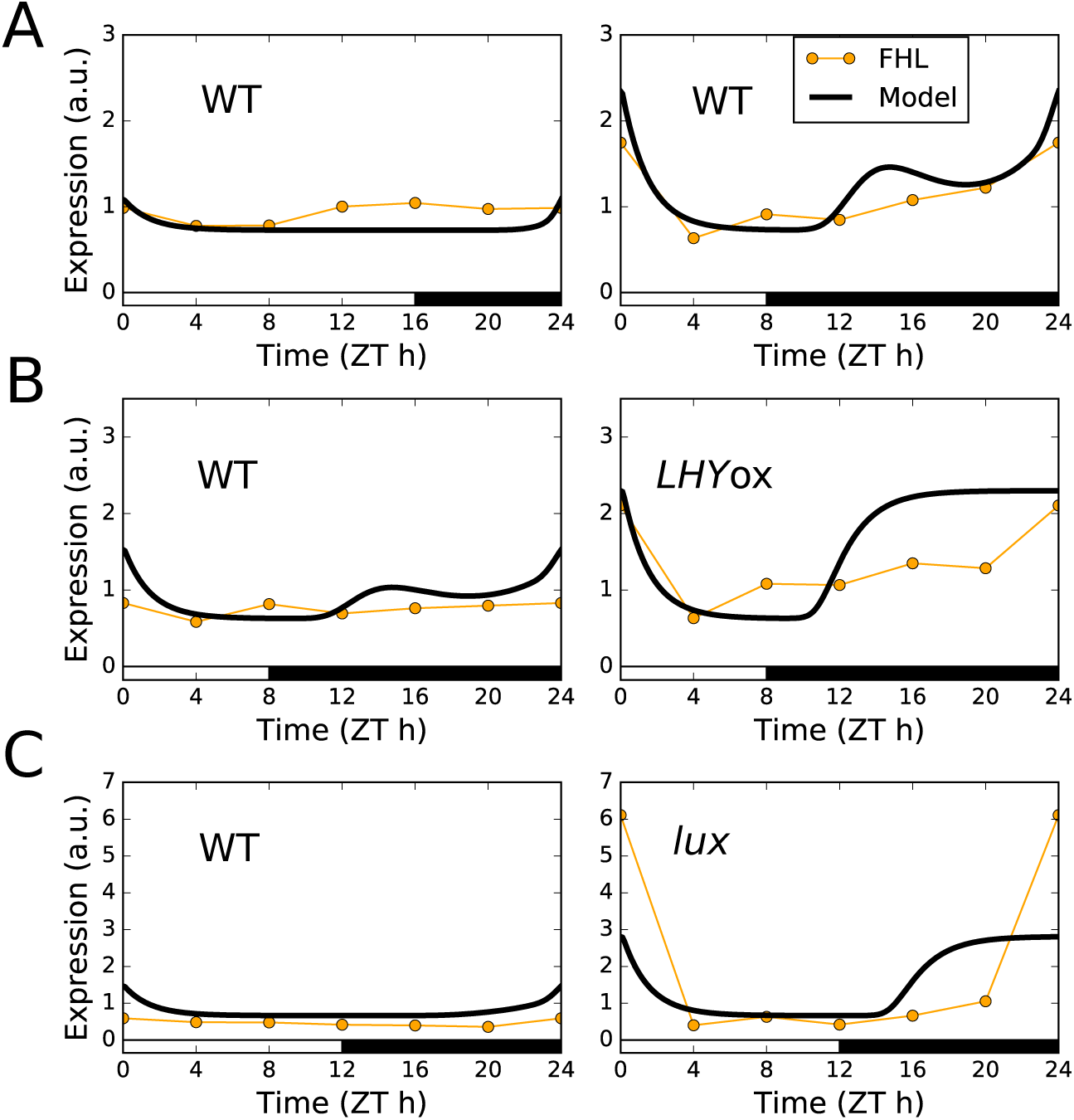
Comparison of model simulations and microarray data for *FHL* expression. (A) WT (Ler) in short compared to long photoperiods. (B) WT (Ler) compared to *LHY*ox in 8L: 16D light/ dark cycles. (C) WT (Col) compared to the *lux* mutant in 12L:12D light/dark cycles.

**Supplementary Figure S4.**
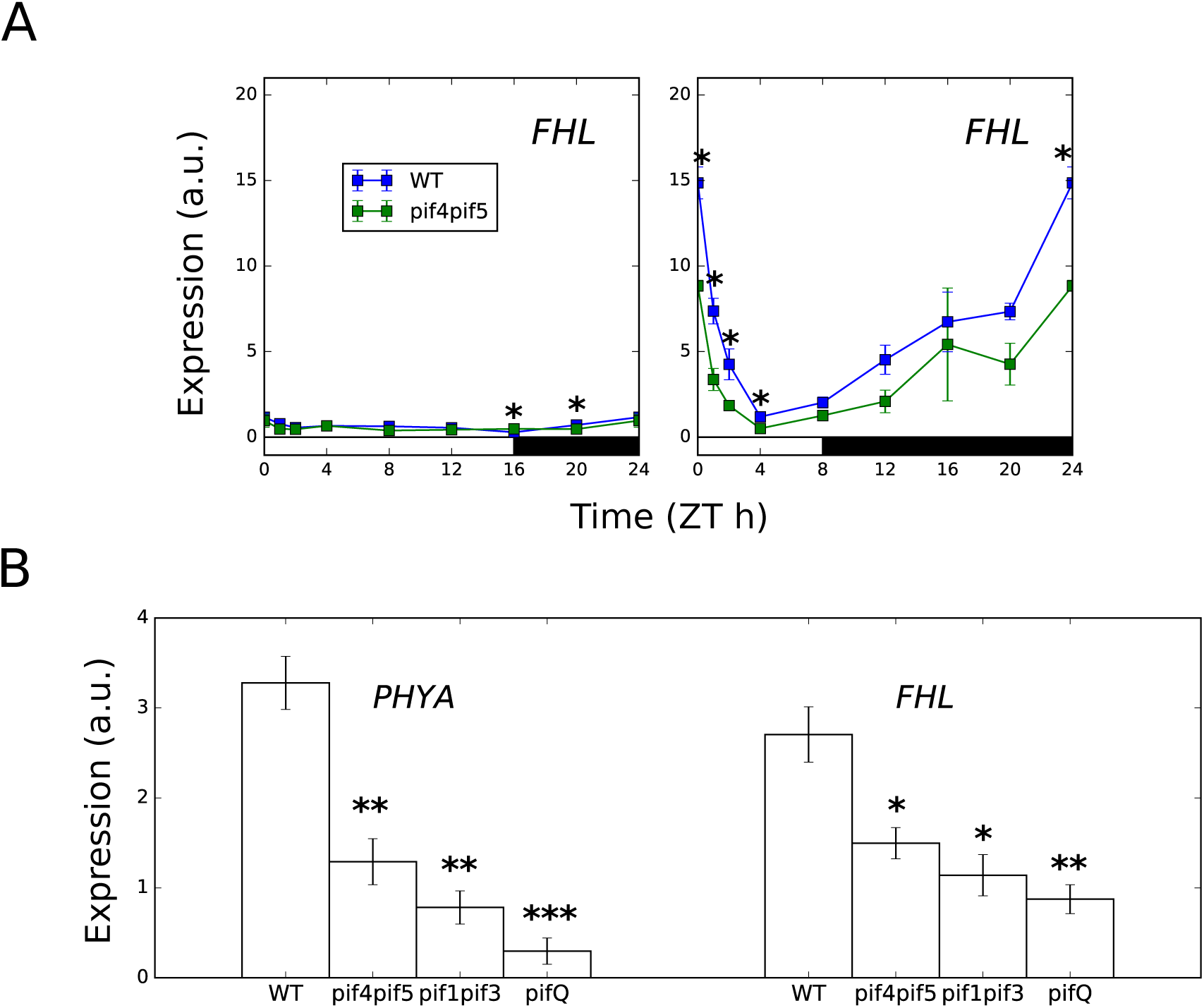
(A) *FHL* expression in short and long photoperiods, in the WT (Col) and the *pif4pif5* mutant. * indicates a difference from WT at p < 0.05 (two-tailed t-test, n=3, error bars represent SEM). (B) *FHL* and *PHYA* expression in short photoperiods at ZT0, in the WT (Col), and the *pif4pif5*, *pif1pif3*, and *pifQ* mutants. Plants were grown for 2 weeks in the stated photoperiod. Expression was measured relative to *ACT7*. *,**,*** indicates a difference from WT at p < 0.05,0.01,0.001 respectively (two-tailed t-test, n=3, error bars represent SEM).

**Figure S5.**
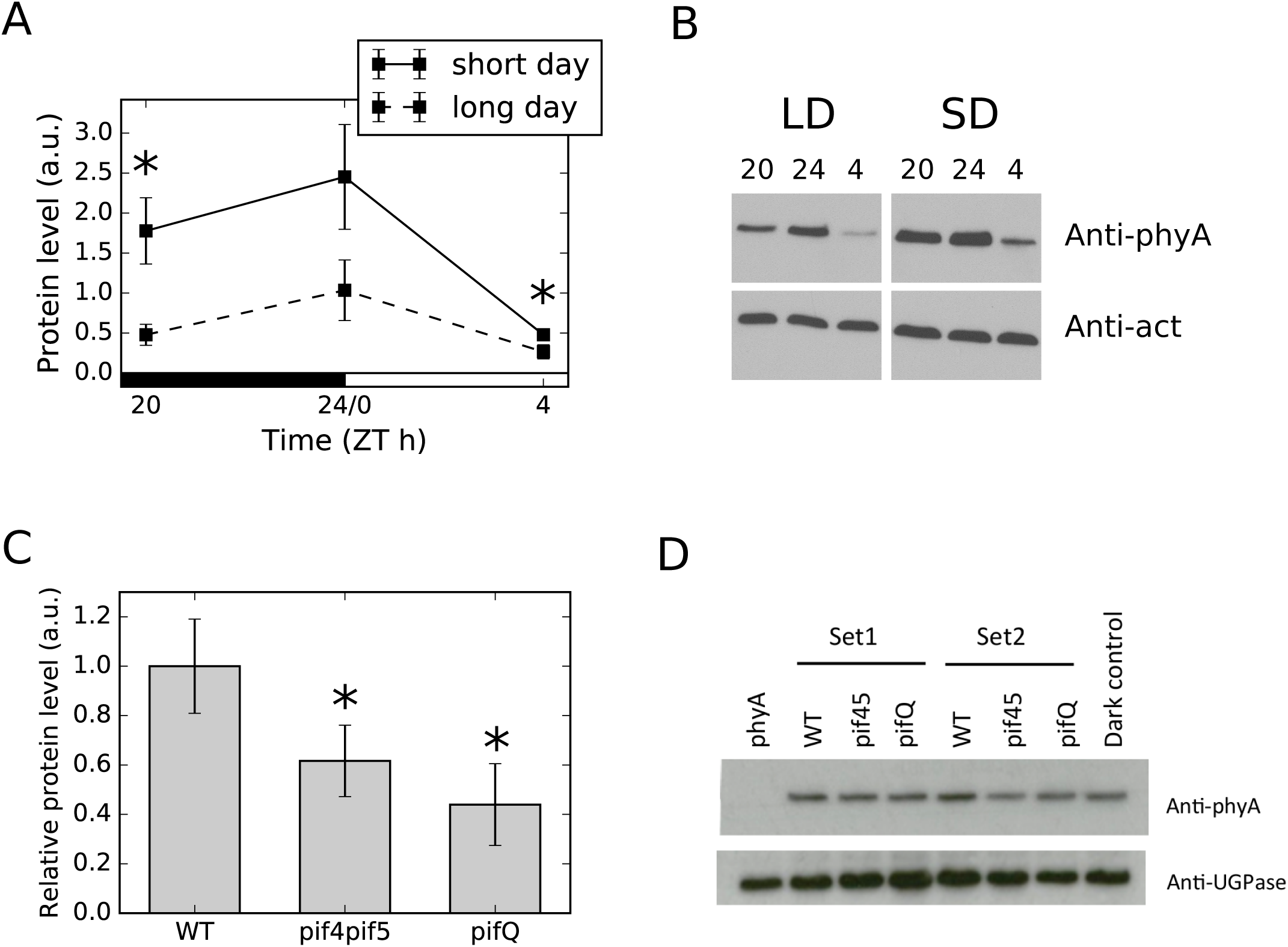
phyA protein quantification. (A) Quantified Western blot data for phyA at three timepoints spanning dawn, in WT (Col), for two-week old plants grown in short and long days, normalised to actin loading standard (error bars represent SEM, n=3, * p<0.05, one-sided t-test). (B) Representative Western blot of data plotted in (A). Note that images shown are taken from the same blot. (C) Quantified Western blot data for phyA at ZT0 (before lights on), in WT (Col) and the pifQ and pif4;pif5 mutants, for plants grown in short days, normalised to a UGPase loading standard (error bars represent SEM, n=4, * p<0.05, one-sided t-test for paired samples). (D) Representative Western blot of data plotted in (C).

**Supplementary Figure S6.**
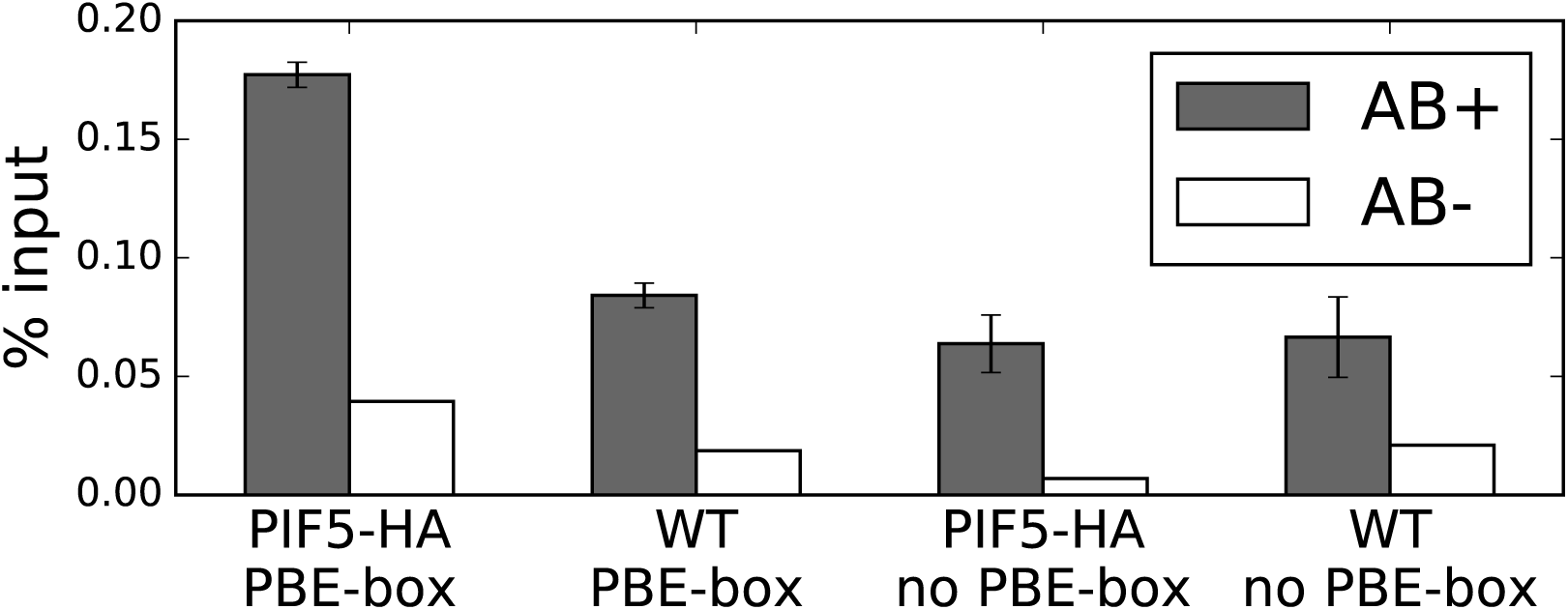
PIF5 ChIP at the *PHYA* promoter. Plants were grown for two weeks in short days (8L:16D white light, 100 µmol/m^2^/s) at 22°C, and samples were collected at the end of the two weeks at ZT0 (n=3, error bars represent SEM).

**Supplementary Figure S7.**
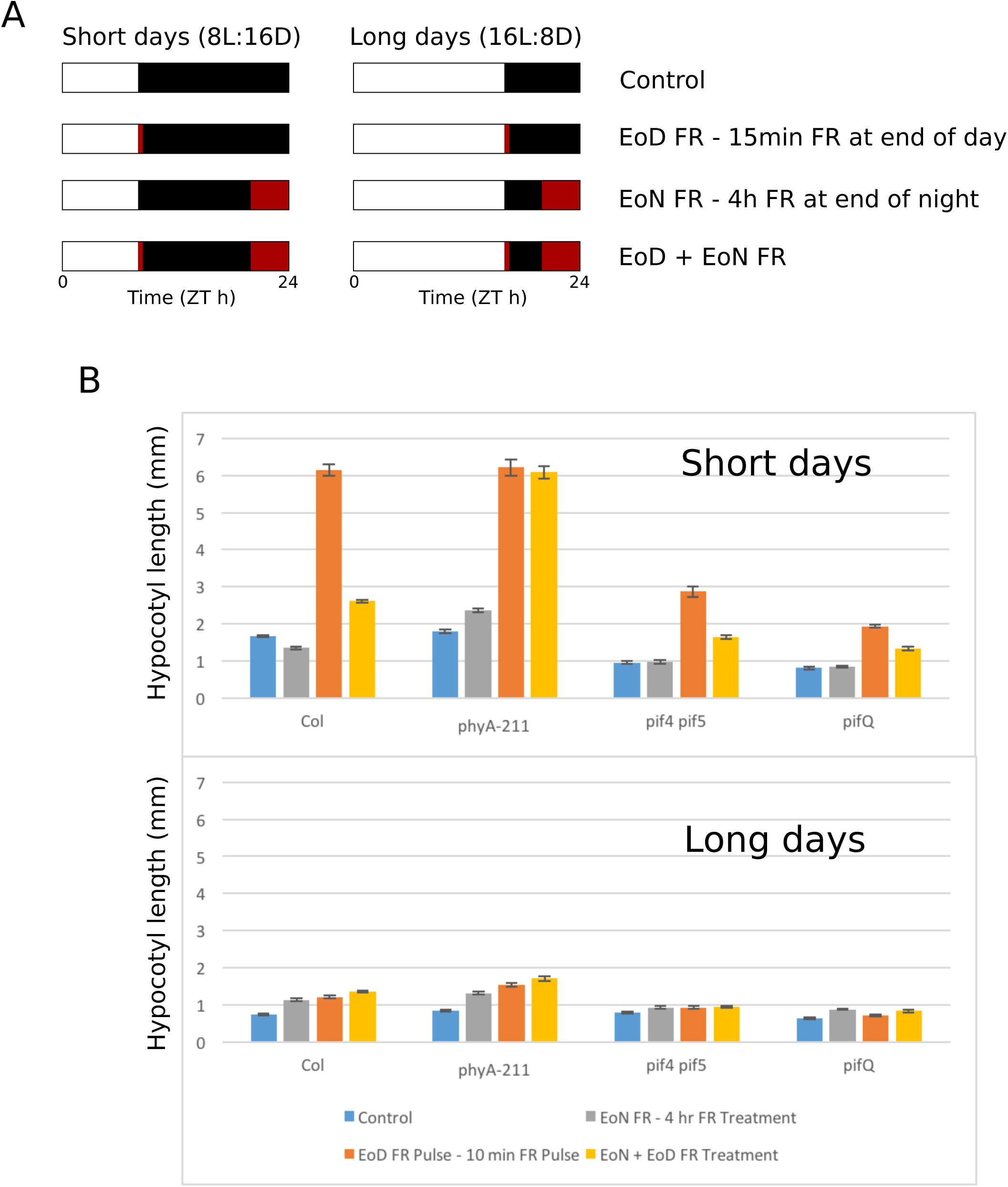
Hypocotyl elongation in response to photoperiod and far-red light. (A) Schematic of light treatments. (B) Hypocotyl measurements for WT (Col0), phyA, pif4;pif5, and pifQ. Plants were grown for 7 days at 22°C in the specified light conditions. Error bars represent SEM. N>12.

**Supplementary Figure S8.**
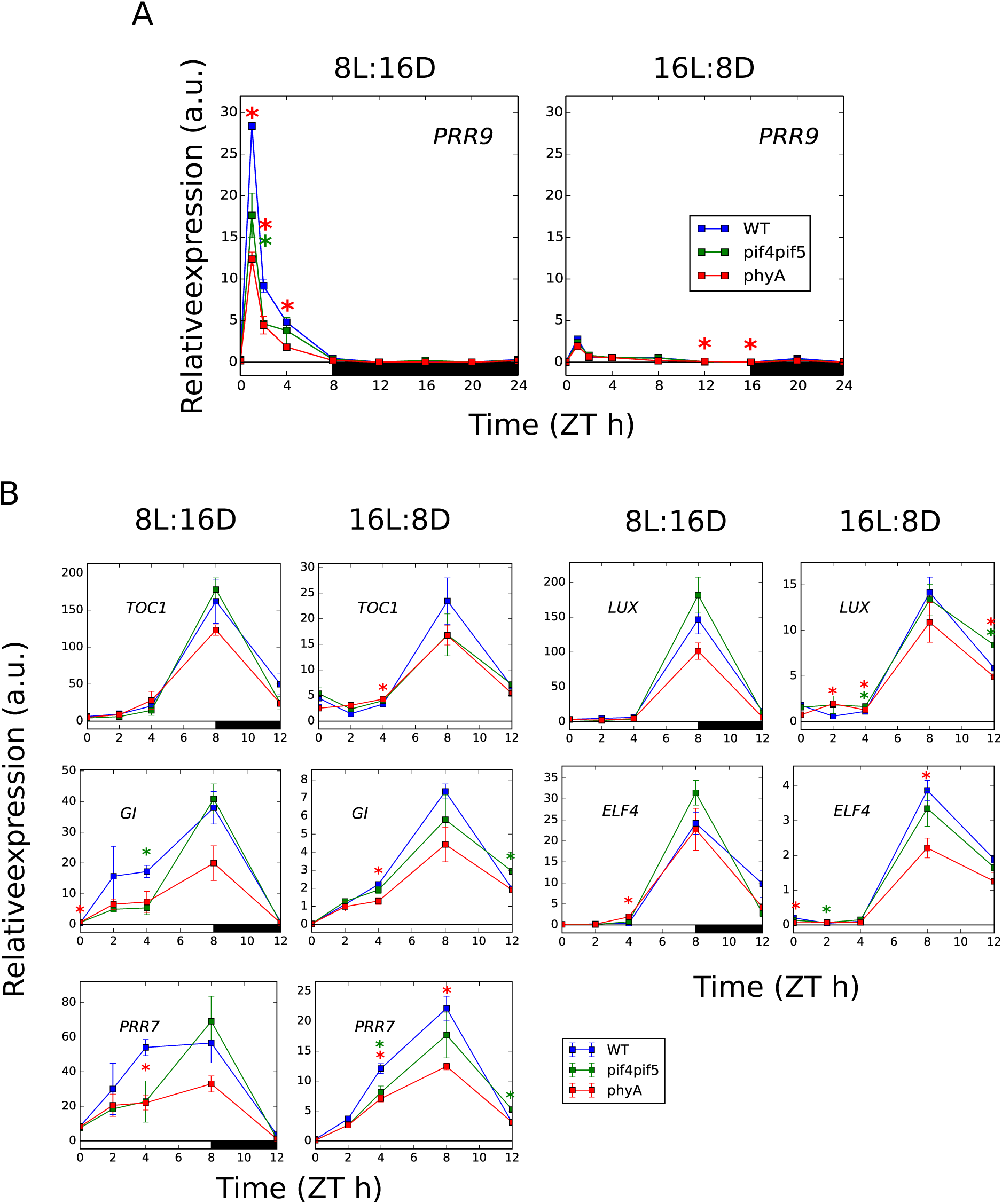
Clock gene expression in response to photoperiod and loss of phyA. (A) qPCR timecourse data for PRR9 in short (left) and long (right) photoperiods, in WT (Col0), pif4;pif5, and phyA (B) qPCR timecourse data for core clock genes at a subset of timepoints between ZT0 and ZT12, in short (left) and long (right) photoperiods, in WT (Col0), pif4;pif5, and phyA. Expression is relative to ACT7. Plants were grown for 2 weeks at 22°C under 100 µmol/m^2^/s white light in the specified photoperiod (n = 3, error bars represent SEM, green and red *s indicates significant difference between WT and pif4;pif5 and phyA, respectively, p < 0.05, two-tailed t-test).

**Supplementary Figure S9.**
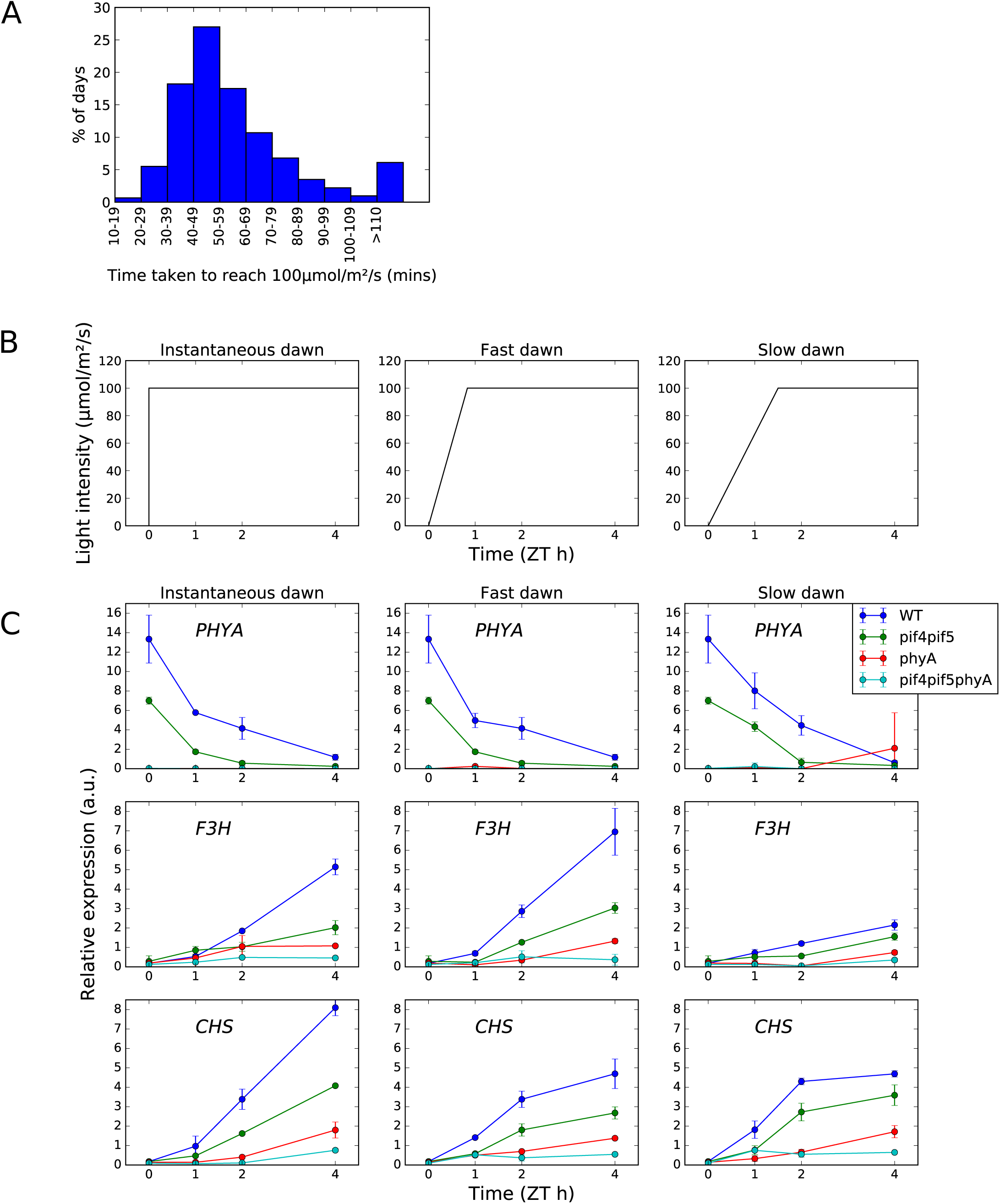
phyA signalling in simulated natural dawn conditions. (A) Histogram of the time taken for the light intensity to reach 100 µmol/m^2^/s on days with short photoperiods in Edinburgh, UK (see Supporting Information for details). (B) Schematic of the experimental protocol to simulate natural dawn based on weather data. (C) qPCR timecourse data for *PHYA*, *F3H*, and *CHS* in the three light conditions shown in (B), in WT (Col0), pif4;pif5, phyA, and pif4;pif5;phyA mutants. Expression is relative to ACT7. Plants were grown for 2 weeks at 22°C under 100 µmol/m2/s white light in the short photoperiods (n = 3, error bars represent SEM).

**Supplementary Figure S10.**
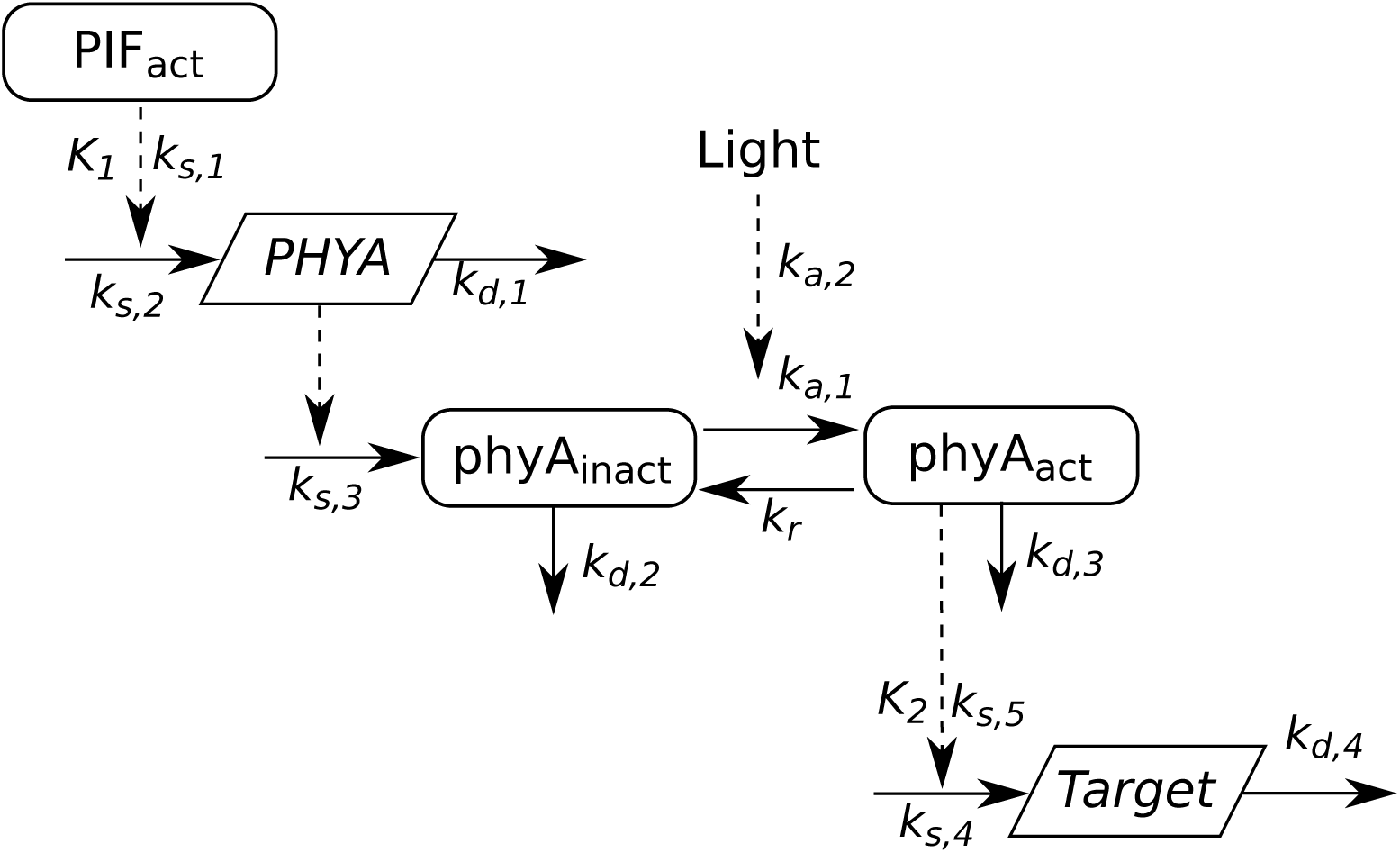
Schematic of phyA model. Rectangles indicate protein species. Trapezoids indicate transcripts. Solid lines indicate mass transfer (synthesis and turnover of molecules, and conversion of phyA between inactive and active forms). Dashed lines indicate regulatory influences.

